# Reconfigurable MRI coil technology can substantially reduce RF heating at the tips of bilateral deep brain stimulation implants

**DOI:** 10.1101/474015

**Authors:** Laleh Golestanirad, Boris Keil, Sean Downs, John Kirsch, Behzad Elahi, Julie Pilitsis, Lawrence L Wald

## Abstract

Patients with deep brain stimulation (DBS) implants can significantly benefit from magnetic resonance imaging (MRI) examination, however, access to MRI is restricted in this patients because of safety concerns due to RF heating of the leads. Recently we introduced a patient-adjustable reconfigurable MRI coil system to reduce the SAR at the tip of deep brain stimulation implants during MRI at 1.5T. A simulation study with realistic models of single (unilateral) DBS leads demonstrated a substantial reduction in the local SAR up to 500-fold could be achieved using the coil system compared to quadrature birdcage coils. Many patients however, have bilateral DBS implants and the question arises whether the rotating coil system can be used in for them. This work reports the results of phantom experiments measuring the temperature rise at the tips of bilateral DBS implants with realistic trajectories extracted from postoperative CT images of 10 patients (20 leads in total). A total of 200 measurements were performed to record temperature rise at the tips of the leads during 2 minutes of scanning with the coil rotated to cover all accessible rotation angles. In all patients, we were able to find an optimum coil rotation angle and reduced the heating of both left and right leads to a level below the heating produced by the body coil. An average heat reduction of 65% was achieved for bilateral leads. Reconfigurable coil technology introduces a promising approach for imaging of patients with DBS implants.

## Introduction

Deep brain stimulation (DBS) is a reversible and adjustable neurostimulation technique in which specific brain structures and circuits are electrically stimulated by means of metallic electrodes connected to an implantable pulse generator (IPG) via subcutaneous insulated wires. The US Food and Drug Administration (FDA) approved DBS as a treatment of essential tremor and Parkinson’s disease (PD) in 1997, dystonia in 2003 [1], obsessive-compulsive disorder (OCD) in 2009 [2], and recently epilepsy in 2018 [3]. Beside this, in recent years DBS has been extensively used in open-label studies to treat Tourette’s Syndrome [4, 5], bipolar disorder [6], Schizophrenia [7], treatment-resistant depression [8, 9], and freezing of gait [10]. The exponential increase in indications of use and application of DBS in neuropsychiatric disorders parallels the large availability and need for MRI. Particularly, MRI has been largely leveraged to understand the underlying mechanisms of action of neuromodulation systems and more recently to guide therapy. The clinical community however, has been cautious in adopting MRI in DBS patients mostly due to concerns about interaction of MRI fields and the implanted device. This includes for example, distortions that affect image quality (e.g., an in-situ device induces susceptibility artifacts), interaction with static magnetic field, loss of device functionality due to interference of gradient fields, but most importantly the potential of thermal injury to patients due to interaction with radiofrequency (RF) fields. Such concerns have led many centers to refrain from performing postoperative MRI on DBS patients mainly to adhere to industry-proposed warnings [11]. In some cases, patients have faced the proposition of explanting their neurostimlator to receive a diagnostic MRI [12]. With the advances in neurostimulation technology, some of the above-mentioned concerns have been mitigated: reduction of ferromagnetic material has reduced the risk of device dislodgment due to static magnetic field [13] and improved device programming has reduced the risk of malfunction due to effect of gradient fields [14]. RF heating however, remains a major issue. The RF safety of MRI in patients with elongated implants has been extensively discussed in literature [15–18]. Techniques based on parallel transmit pulse tailoring [19–25] and surgical lead management [26, 27] have shown promising results in reducing the temperature rise near implanted wires in phantoms during MRI at 1.5T and 3T, but such techniques have not been implemented in clinical settings yet. Recently we introduced a patient-adjustable reconfigurable MRI coil with promising results in reducing SAR amplification near tips of implanted leads [28, 29]. The coil system consisted of a linearly-polarized rotating birdcage transmitter and a 32-channel close-fit receive array. Such a linearly-polarized transmit coil has a slab-like region of low electric field which can be steered to coincide with the implant trajectory by rotating the coil around patient’s head. This will significantly reduce the coupling of electric fields with implanted wires, which in turn reduces the RF induced currents on the leads and the SAR at the tip. In a simulation study with patient-derived realistic models of 9 unilateral DBS leads we demonstrated that a substantial reduction in the local energy deposition can be achieved using this technique, reducing the SAR by up to 500-fold compared to quadrature birdcage coils [29]. In practice however, most patients with PD or essential tremor receive bilateral implants as it has been shown more effective at controlling appendicular and midline tremor [30]. Additionally, 52% of patients who originally receive unilateral DBS eventually need bilateral stimulation [31]. Considering the fact that the optimal coil rotation angle to minimize the local SAR is dependent on the specific lead trajectory [29], the question arises as whether or not the rotating coil system can ever be used in patients with bilateral DBS implants. In other words, as the maximum SAR-reduction occurs when the lead is fully contained in the low E field region of the coil, it is crucial to assess if there exists a rotation angle that reduces the heating of both left and right DBS leads to the level below the heating generated by a conventional quadrature coil.

In this work we report, for the first time, measurement results of temperature rise at the tips bilateral DBS lead implants with realistic trajectories during MRI at 1.5 T using the rotating coil system. Implant trajectories were extracted from CT images of 10 patients with bilateral DBS leads (20 leads in total). Anthropomorphic head phantoms were constructed and implanted with bilateral lead wires and positioned inside the coil system with temperature probes attached at their tips. For each experiment, the coil was rotated around the head phantom covering the full range of accessible angles and the temperature at the tips of implants was recorded at each rotation angle during 2 minutes of continues RF exposure. A total of 200 measurements were performed (10 patients×16-22 measurements per patient). We found that for all realistic bilateral lead trajectories, there existed an optimum coil position that substantially reduced the heating of both left and right DBS leads compared to the scanner body coil. An average heat reduction of 65% was achieved in the case of bilateral leads. In the case of single leads, an average heat reduction of 80% was achievable using the rotating coil system.

This work reports the first experimental results of successful application of the reconfigurable coil technology to reduce heating of bilateral DBS leads. The results produce highly relevant data for regulatory guidelines and premarket applications.

## Methods

### The reconfigurable coil system

Details of the theory and construction of the reconfigurable birdcage coil system are given in our previous works [28, 29]. In brief, the coil consists of a 16-rung linearly-polarized low-pass birdcage transmitter and a 32-channel close-fit receive array (Figure *1*A). Such a linearly-polarized birdcage has a slab-like region of low electric field which can be steered by rotating the coil around patient’s head to coincide with the implant trajectory. Figure 1B shows the distribution of coil’s electric field on a central axial plane for two different coil rotation angles (Figure from refs [28, 29]). A generalized implant trajectory has been also depicted for visualization (implant was not included in simulations that generated field distributions). As it can be observed, by rotating the coil around patient’s head and positioning it at an optimal angle, a substantial portion of the lead trajectory can be contained within the low E field region. This, reduces the coupling of the electric fields and conductive wires which in turn reduces the SAR at the implant’s tip. Figure 1C shows the distribution of 1g-averaged SAR at different coil rotation angles for a generic implant trajectory (Figure from ref [28, 29]). In theory, a SAR-reduction factor of up to 500-fold is achievable with such a coil for generic and realistic unilateral DBS trajectories [28, 29]. It is important to note however, that the SAR-reduction performance of the coil and the optimum rotation angle that minimizes the SAR at the tip of each implant is dependent on the lead trajectory [29]. This is in line with previous studies that have emphasize on the importance of the relationship between implant’s geometry, and the phase and orientation of incident electric field of MRI transmit coils [21, 32-37].

**Figure 1:**
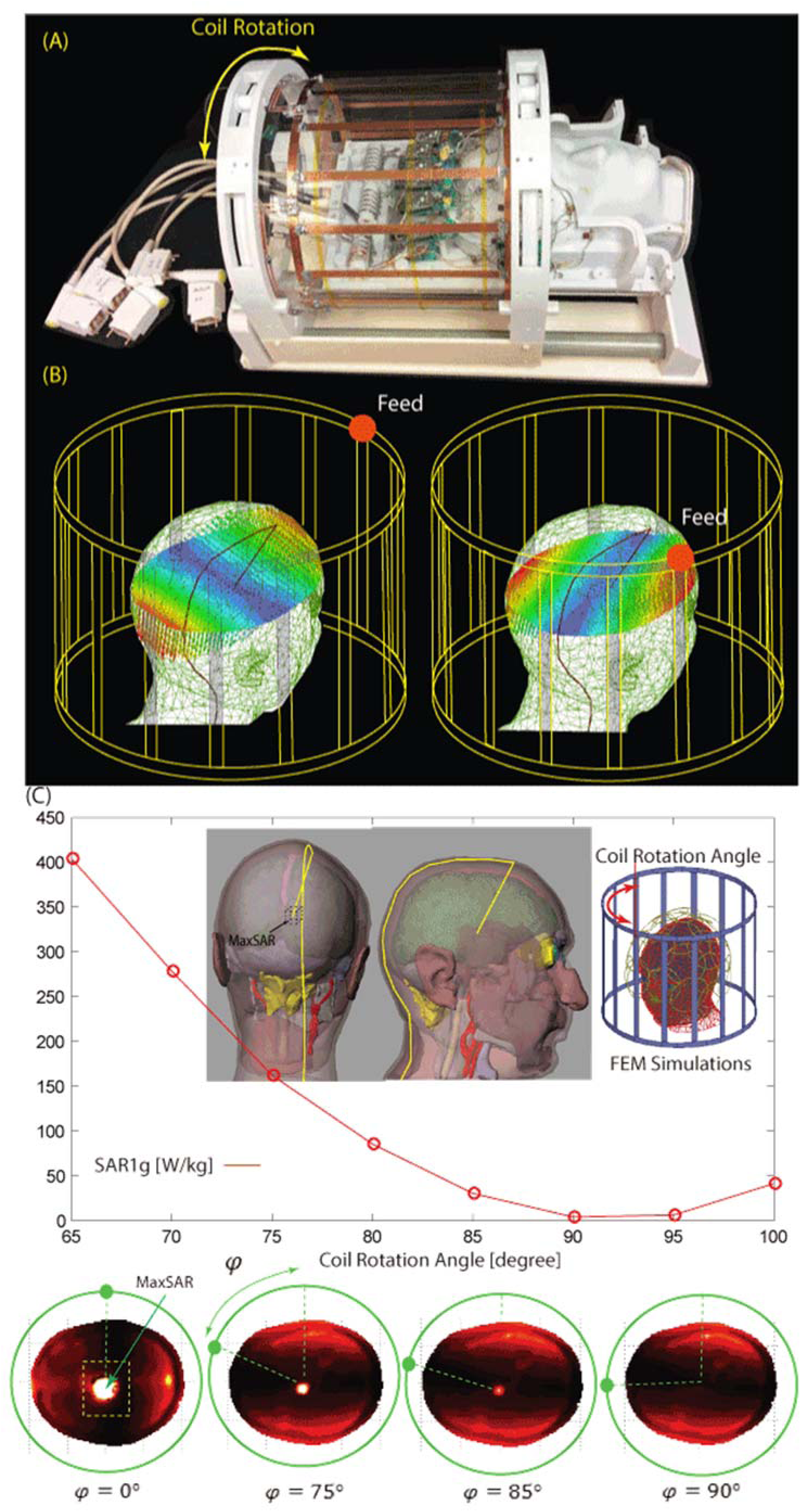
An overview of the reconfigurable MRI coil system. (A-B) View of constructed prototype and its simulated incident electric field profile (C) Calculated 1g-averaged SAR at the tip of a generic DBS implant for different coil rotation angles. The SAR is minimum when the implant is maximally contained in the low E-field region of the coil.

Figure *2*B-C shows the constructed coil prototype on the patient table and the view from back of the coil. The mechanical housing allows the transmitter to rotate smoothly around the receive array and be locked in pre-marked positions which are 5*°* apart. When positioned at the magnet’s iso-center, the coil can be accessed and rotated from the back of the bore, eliminating the need to take the patient out for adjustments. Due to mechanical restrictions imposed by positioning of the cables however, the coil cannot rotate in a full circle. The range of accessible angles (0*°*-140*°* and 220*°*-360*°*) are annotated in Figure *2*.

**Figure 2:**
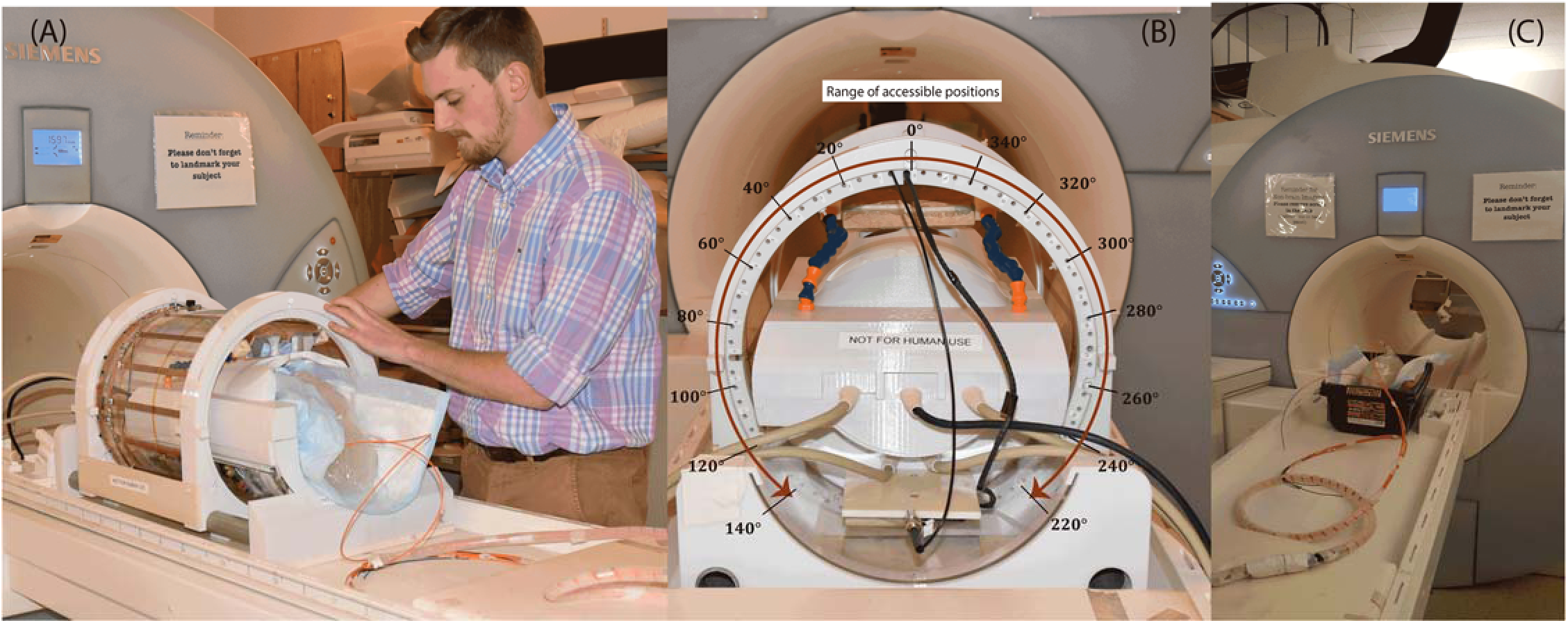
(A-B) The positioning of the rotating coil system on the patient’s table and the range of accessible rotating angles. (C) Phantom scanned with the body coil for comparison. The body coil generated the same level of whole-head SAR as the rotating coil.

## DBS leads and head phantoms

Postoperative CT images of ten patients who had undergone surgery for bilateral DBS implantation was used to extract lead trajectories (Figure *3*). A total of 20 leads were constructed for measurements. Amira (Thermo Fisher Scientific, Waltham MA) was used for image segmentation and construction of the preliminary 3D surface of the leads. First, a thresholding mask was applied to select the hyper dense DBS lead from CT images using Amira’s segmentation module (Figure *4* A-B). Threshold values were selected manually on a case-by-case basis such that the resulting mask produced a smooth continuous 3D surface. 3D lead surfaces were exported to a CAD tool (Rhino3D®, Robert McNeal and Associates, Seattle, WA) in which lead trajectory lines were manually extracted, thickened (4mm diameter), and prepared for 3D printing (Figure *4* C). Lead guides were then 3D printed out of polycarbonate plastic using a Fortus 360mc 3D printer (Stratasys, Eden Prairie, MN, USA). Two pieces of insulated wire (Ga 14, 40 cm long, 1cm exposed tip) were shaped around 3D printed guides to follow the left and right lead trajectories (Figure *4* D). Wires were rigid enough to maintain their shape once they were routed around the plastic guides and were detached from the guide before being implanted into the head phantom.

An anthropomorphic head phantom was designed and 3D-printed based on the structural MRI of a healthy volunteer. The mold was composed of two sagittal sections connected through a rim (Figure *5* A). Phantom dimensions were approximately 16 cm ear-to-ear and 27 cm from top of head to bottom of neck. The mold was filled with agarose-doped saline solution (5L water, 14g NaCl) through a hole at the bottom. A relatively high percentage of agarose (4%) was used which resulted in a semi-solid gel that could stand alone and support the implants. The electric properties of the gel was measured using a dielectric probe kit (85070E, Agilent Technologies, Santa Clara, CA) and a network analyzer) to be *τ = 0.47 S/m* and *ε_r_ = 77.* Leads were implanted into the gel following the entrance point, angle, and trajectories as observed from CT images of the patient (Figure *5* B). Fluoroptic temperature probes (OSENSA, BC, Canada) were secured at the exposed tips of the wires for temperature measurements. A third probe was inserted to the center of the head phantom for background measurements.

**Figure 3:**
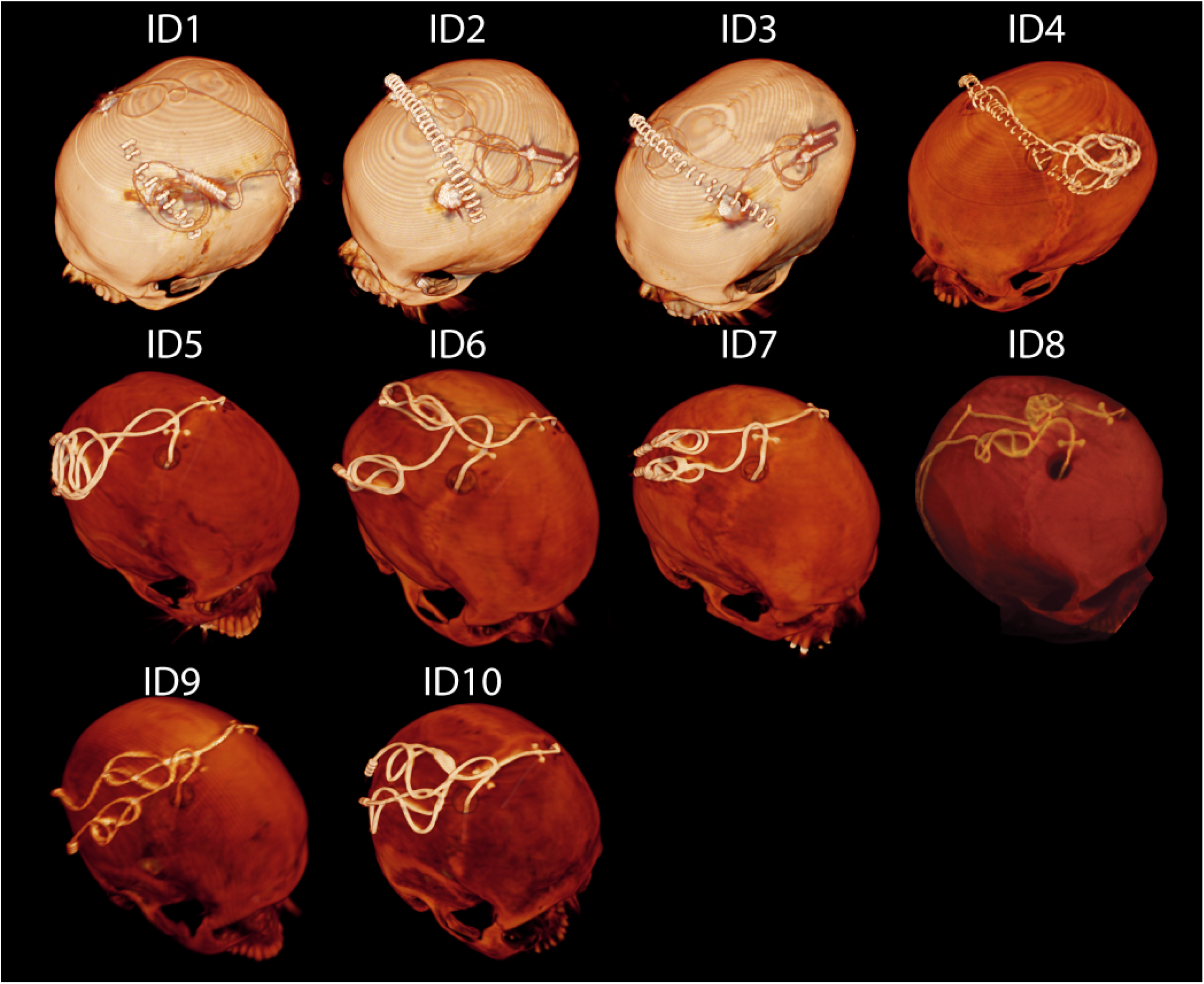
CT images of 10 patients operated for bilateral DBS implantation. For patients 1-4, the IPG was planned to be implanted in the left pectoral pocket. Note that end segments of both leads are routed toward the left side of the head in these patients. In this cases, the left lead was labeled as ipsilateral and right lead was labeled contralateral when representing the temperature results in Figures 6 and 7. For patients 5-10 the IPG was planned to be implanted in the right pectoral region, thus in these patients the right lead was labeled ipsilateral and left lead was labeled as contralateral.

## RF exposure

Experiments were performed at a 1.5T Magnetom Avanto system using the rotating coils system as well as the built-in body coil for comparison. To better control the RF exposure, gradient coils were disabled and a train of 1ms rectangular RF pulses were transmitted using the “*rf_pulse*” sequence from Siemens Service Sequence directory. Scan duration was set to 120 s for both body coil and the rotating coil. The flip angle, however, was adjusted to make sure that all cases had the same level of power absorbed in the head phantom to allow for a fair comparison. The absorbed power in the phantom was determined by subtracting the power dissipated in the coil and the reflected power from the forward power in the coil:

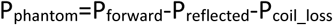

These values, registered as “*TALES powers*” in the scanner log file, are measured in real time at a hardware component called Transmit Antenna Level Sensor (TALES) which resides between the output of the RF amplifier and the coil. In our case, an rf_pulse sequence with TR=6.5 ms and flip angle=180*°* generated 99 W power at the head phantom when transmitted with the body coil. For the rotating coil, the TR was kept the same but flip angle was adjusted to 150*°* which generated 101 W power at the head phantom. Note that considering the weight of the phantom (~4kg), these values generated a global head SAR of ~25W/kg which is significantly higher than the FDA recommended limit of 3.2 W/kg for clinical applications. Such high SAR value was necessary to produce enough heating at the tips of the lead wires to be above the noise in all experiments.

## Temperature measurements

At the start of each experiment, when the head phantom was at iso-center of the body coil, we measured the temperature rise at the tips of right and left implants by transmitting two-minute rf_pulse sequences using the body coil. Left and right leads were labeled as contralateral or ipsilateral depending on the IPG side. For patients 1-4, the IPG was planned to be implanted in the left pectoral pocket (see Figure 2, showing end segments of both leads routed toward the left side of the head). For these patients, the left lead was labeled as ipsilateral and right lead was labeled contralateral. For patients 5-10 the IPG was planned to be implanted in the right pectoral region, thus in these patients the right lead was labeled ipsilateral and left lead was labeled as contralateral. After measurements with the body coil, the phantom was left for 15 minutes to cool down. Measurements with the rotating coil started with the coil at its default position (feed up, *0 = 0°*). The rf_pulse protocol was transmitted for 2 minutes and temperatures at the tips of ipsilateral and contralateral leads as well as the background probe were recorded. The coil was then rotated to the left at 20*°* increments until all accessible rotation angles from 0*°* to 140*°* were covered (see Figure *2*). The coil was consequently repositioned at 0*°* and rotated to the other direction to cover angles from 360*°*-220*°*. Depending on the heating, the phantom was left for 15-30 minutes between each experiment to return to baseline temperature. At the end of each experiment we added a few extra measurement points around the position that produced the minimal heating on each lead to better estimate the optimum position angle.

**Figure 4:**
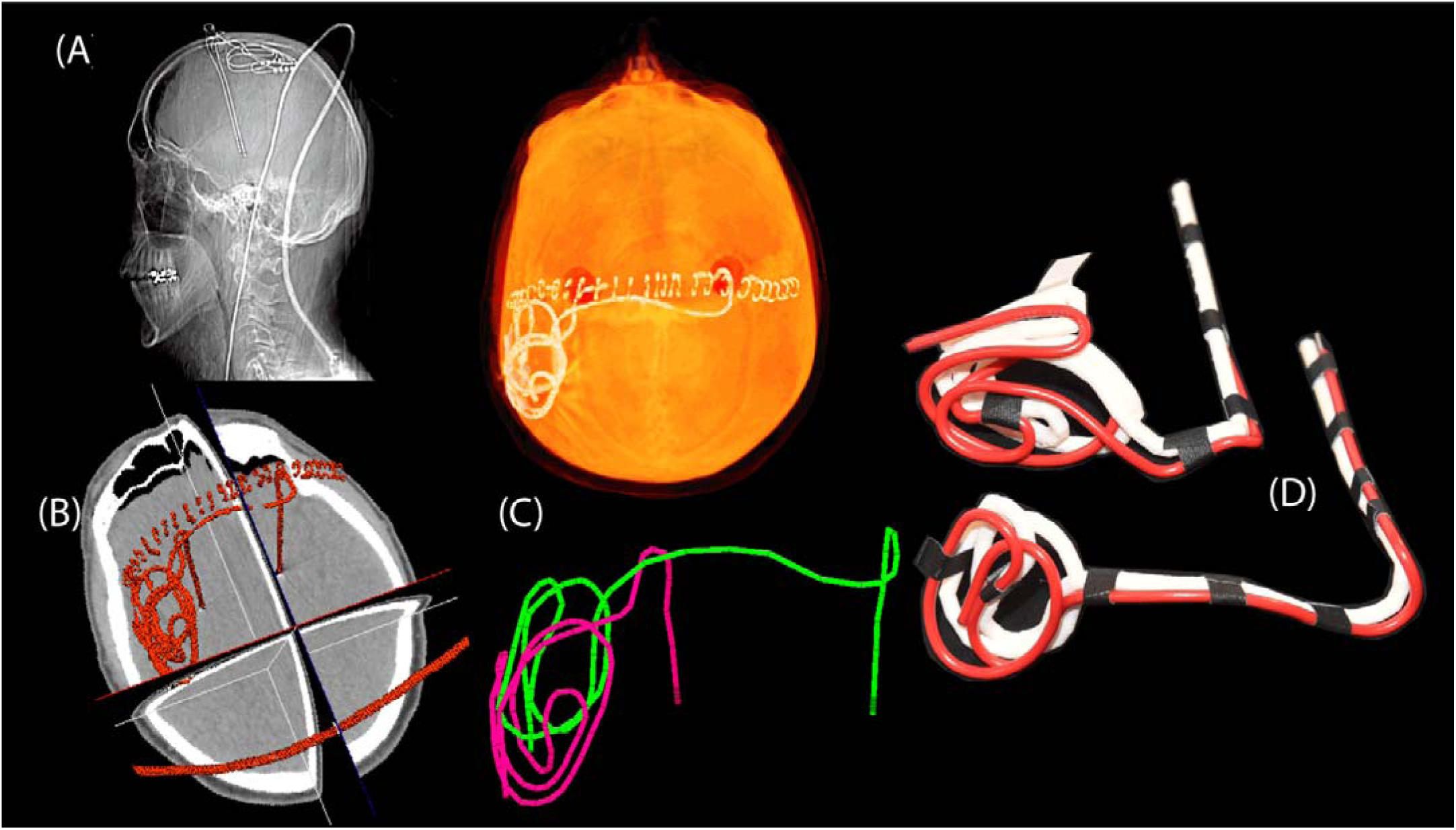
Steps of image segmentation (A-B), 3D model construction (C), and 3D printed plastic guides (D).

**Figure 5:**
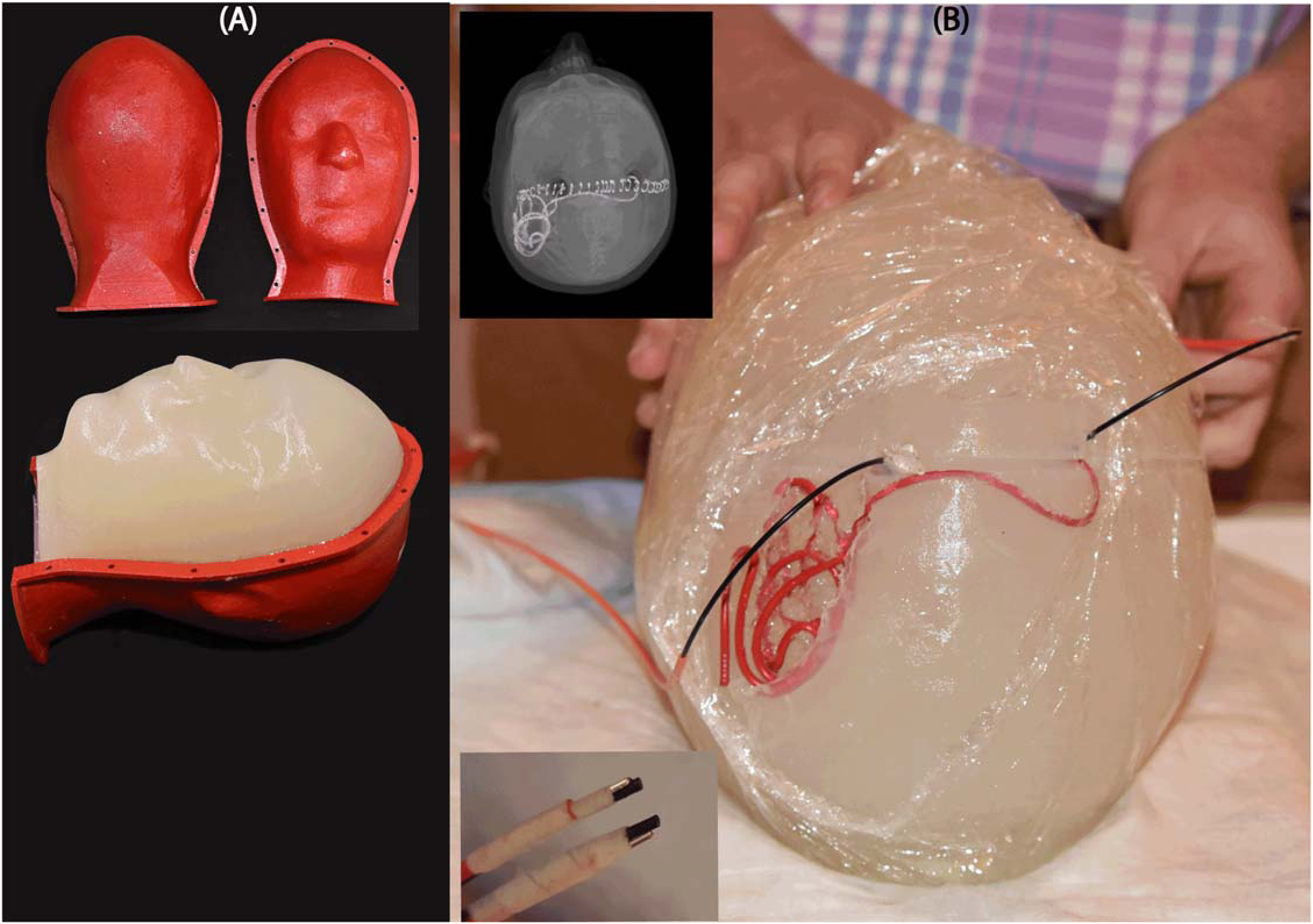
(A) Anthropomorphic head phantom based on MRI of a healthy volunteer (B) 3D printed DBS lead phantoms were used as a guide to shape wires in the form of patient-derived trajectories. Wires were implanted into semi-solid anthropomorphic head phantoms with temperature probes attached to their tips for temperature measurement during MRI with the body coil and the rotating coil system.

## Results

Figure 6 and Figure 7 show the result of temperature measurements at the tips of ipsilateral and contralateral leads of patients 1-10 for the range of accessible coil rotation angles during 2-minute RF exposure at the global SAR level of 25 W/kg. For each patient, the temperature rise at the tips of the leads are also given for body coil that generated the same global SAR (straight dashed lines).

**Figure 6:**
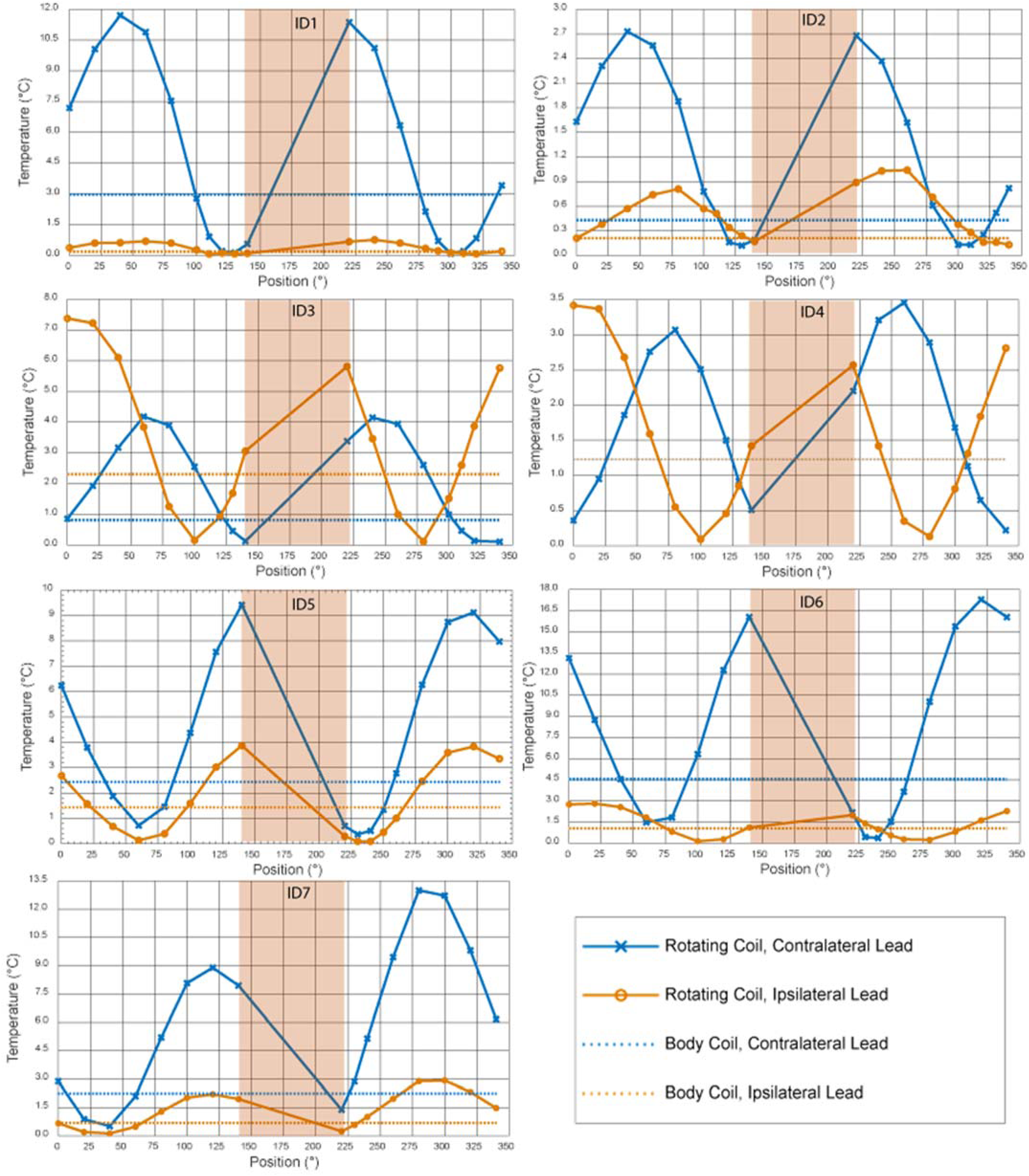
Temperature rise at the tips of ipsilateral and contralateral leads after 2 minutes of RF exposure. The highlighted area is the range of non-accesible coil angles.

**Figure 7:**
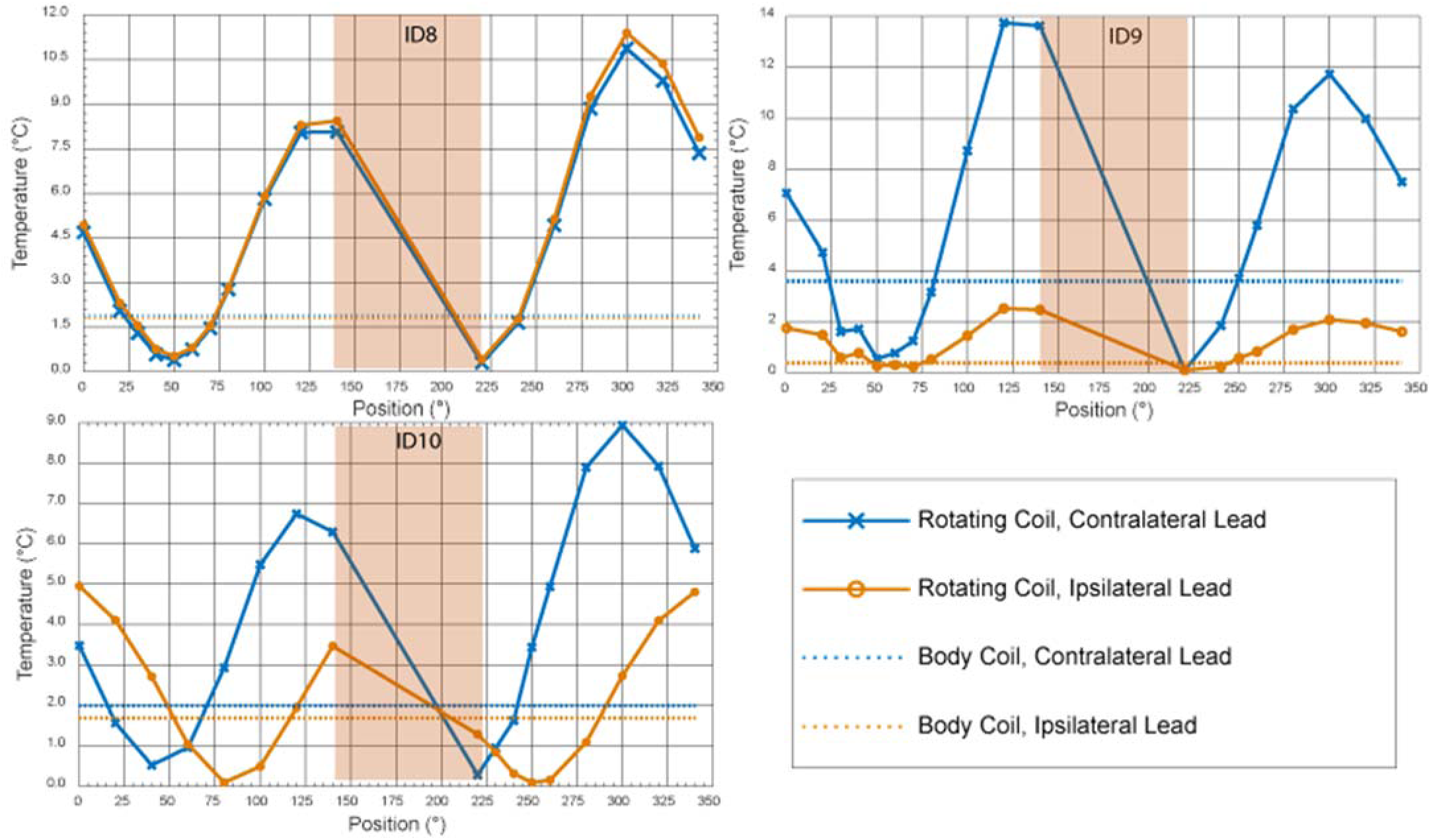
Temperature rise at the tips of ipsilateral and contralateral leads after 2 minutes of RF exposure (continued). The highlighted area is the range of non-accesible coil angles.

To quantify the performance of the reconfigurable coil to reduce the heating of implanted leads, we defined a factor called *heat reduction percentage* or HRP for each lead and at each coil rotation angle as:

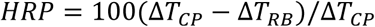

where *∆T_CP_* is the temperature rise at the tip of the lead at the end of the 2-minute measurement with the built-in body coil and *∆T_RB_* is the temperature rise at the end of 2-minute measurement with the rotating birdcage coil. A positive HRP at a rotation angle *θ* indicates that for the same level of global SAR, the rotating birdcage coil positioned at *θ* will generate less heating at the tip of the implanted wire compared to the body coil. In contrast, a negative HRP at an angle *θ* indicates that the rotating coil positioned at *θ* generates more heating at the tip of the implanted wire compared to the body coil when both coils generate the same level of global SAR.

### Optimal rotation angle for single leads

Table 1 gives the HRP values for ipsilateral and contralateral leads of each patient at two distinct coil rotation angles which minimized either the heating of ipsilateral lead or the heating of contralateral lead. Table entries highlighted green show the lead for which the heating was minimized. For example, in patient 9, coil at *θ = 230°* minimized the heating of ipsilateral lead (HRP_Ipsi_=97) and coil at *θ = 40°* minimized the heating of contralateral lead (HRP_contra_=63). As it can be observed from Table 1 (as well as Figures 7 and 8), for each of the 20 leads there existed an optimum coil rotation angle that reduced the heating of the lead to a level well below the heating produced by the body coil. On average, a substantial heat reduction of 80%±19% was achieved for single leads. In 33% of cases (patients 5,7 and 8) the optimum rotation angles that minimized the heating of ipsilateral and contralateral leads were the same. For the rest of patients, the optimal angles for ipsilateral and contralateral leads were different and in 40% of cases (patients 2,3,4, and 10) the rotation angle that minimized the heating at one lead increased the heating at the other one.

**Table 1:** HRP values for ipsilateral and bilateral leads of patients 1-10. For each patient, HPR is given for rotating angles that minimized the heating either at ipsilateral or contralateral lead

| | | $HRP = 100(\Delta T_{CP} - \Delta T_{RB})/\Delta T_{CP}$ | |
| --- | --- | --- | --- |
| Patients | Coil angle | Ipsilateral Lead | Contralateral Lead |
| ID1 | 130° | 61 | 96 |
|  | 300° | 33 | 97 |
| ID2 | 340° | 19 | -91 |
|  | 300° | -81 | 70 |
| ID3 | 280° | 95 | -218 |
|  | 340° | -149 | 87 |
| ID4 | 100° | 93 | -104 |
|  | 340° | -128 | 82 |
| ID5 | 230° | 93 | 83 |
| ID6 | 100° | 87 | -39 |
|  | 240° | 7 | 90 |
| ID7 | 40° | 82 | 77 |
| ID8 | 220° | 77 | 84 |
| ID9 | 230° | 97 | 19 |
|  | 40° | 75 | 63 |
| ID10 | 250° | 95 | -73 |
|  | 40° | -60 | 74 |

### Optimal rotation angle for double leads

From the results of the previous section one can see that for most realistic bilateral DBS implants there is not an optimal coil rotation angle that maximally reduces the heating of both ipsilateral and contralateral leads. For all patients however, it was possible to find an intermediate coil position that reduced the heating of both leads to a level below the heating produced by a conventional body coil, albeit this position was not always optimum for each lead alone. Table 2 shows the rotation angle that maximized the value of HRP_total_=HRP_ipsi_+HRP_contra_. As it can be observed from the table, for all ten patients we were able to reduce the heating of both ipsilateral and contralateral leads to a level below the heating produced by the body coil. When optimized for bilateral leads, an average heat reduction factor of 65%±25% was achieved.

**Table 2:** HRP values for ipsilateral and bilateral leads of patients 1-10. For each patient, HPR values are given for the rotating angle that maximized HRP_ipsi_+HRP_contra_.

| Patients | Coil angle | HRP=100($\Delta T_{CP} - \Delta T_{RB}$)/$\Delta T_{CP}$ | |
| --- | --- | --- | --- |
|  |  | Ipsilateral Lead | Contralateral Lead |
| ID1 | 130° | 61 | 96 |
| ID2 | 140° | 19 | 56 |
| ID3 | 130° | 27 | 44 |
| ID4 | 130° | 63 | 44 |
| ID5 | 230° | 93 | 83 |
| ID6 | 250° | 48 | 66 |
| ID7 | 40° | 82 | 77 |
| ID8 | 220° | 77 | 84 |
| ID9 | 220° | 72 | 97 |
| ID10 | 220° | 24 | 86 |

## Discussion and Conclusion

Patients with DBS implants can significantly benefit from MRI examinations. MRI is preferred to computed tomography (CT) scans because of its non-ionizing nature and the vast range of available sequences for investigating both anatomy and function of the brain. There are several circumstances in which patients with DBS implants may need MRI. Firstly, most patients require brain MRI as a part of diagnostic workup for various comorbidities including postoperative hemorrhage, ischemic stroke, brain tumors, or other neurological and musculoskeletal conditions such as osteoarthritis and suspected stroke [38, 39]. In fact, the American College of Radiology has recognized such conditions as the “criteria for diagnostic MRI with no equivalent test holding a similar rating” [40]. Secondly, the non-ionizing nature and superior soft tissue contrast of MRI has made it highly preferable for postoperative electrode localization. The precise evaluation of implanted electrodes’ location is essential to analyze differential effects of stimulation on specific symptoms and evocation of adverse events. This is particularly important considering that the accuracy of implantation, defined as the difference between intended (or planned) and the actual (or implanted) electrode coordinates play an important role in determining the outcome [41, 42]. Current neuroimaging techniques for DBS electrode localization are mostly based on the co-registration of post-operative CT images and preoperative MRI, which has limited accuracy and warrants cumbersome post-processing [43, 44]. Thus, postoperative application of MRI for electrode localization is highly desirable. The third scenario is the application of functional MRI to investigate the neuromodulatory effect of DBS and to determine the underlying functional architecture of affected brain networks. Recent insights into connectivity-based models of brain function have transformed our understanding of underlying mechanisms of DBS [45, 46] and such frameworks will be critical for evaluation of therapeutic modulation hypothesis in the future. Finally, intraoperative MRI is being increasingly used for stereotactic surgery because of its superb anatomical resolution, and the trend is likely to accelerated with the advancements in MRI hardware technology.

Until very recently, Medtronic (Medtronic Inc., Minneapolis MN) was the only manufacturer of MR-conditional DBS systems [47]. In July 2018 Abbott (Abbott Laboratories, Chicago, IL) received FDA approval for their MR-conditional DBS system. Both manufactures’ guidelines however impose restrictive conditions for imaging: only 1.5 T horizontal systems are allowed, and only pulse sequences with specific absorption rate that is significantly lower than the FDA recommended level for global head SAR are permitted. Precisely, Medtronic system recommend 0.1 W/kg whole-head SAR for all types of transmit coils, and Abbot recommends 0.1 W/kg when using body coils for transmit and 0.8 W/kg when using local head transmit coils, whereas the FDA recommended SAR limit in the absence of implants is 3.2 W/kg. Such limitations are problematic as MRI protocols that optimally image DBS leads and subcortical structures tend to have much higher SARs than current recommendations allow [48]. Developing novel MRI technologies that allow for safe imaging of patients with DBS implants in situ is thus, highly desirable. If successful, such technology will have a substantial disruptive effect on the field of clinical brain mapping. This, is particularly timely as researchers have just begun to investigate fundamental mechanism(s) by which DBS modulates brain networks. With new therapeutic applications for DBS emerging, there is a vital need for imaging technologies that overcome present limitations and bring MRI to bear on open questions relating to DBS targeting and mechanisms of action.

### Patient-reconfigurable MRI technology: practicality vs. versatility

The idea of generating and steering electric-field free regions in MR transmit coils to reduce implant heating was originally introduced by Atalar’s group [21, 32] and later adopted in studies suggesting design of implant-friendly modes in parallel transmit volume coils at 3T [19, 20, 22, 49]. Techniques based on tailoring the transmit pulse usually adopt a cumbersome simulation approach to determine the magnitude and phase of the signal at each transmit channel to shape the electric field in the region of interest (i.e., around the implant) while maintaining a user-defined threshold of B_1_ homogeneity throughout the sample. Although using a multi-channel transmit methodology allows more degrees of freedom for field tailoring, and thus could potentially achieve a better SAR reduction for complicated implant trajectories (e.g., multiple leads), such benefit comes with the drawback of rendering the technique complicated for everyday applications in clinic. The rotating coil system introduced here has the advantage of having a very simple setup, however such ease of operation comes at the expense of lack of control on the shape of the low electric field region. In other words, apart from *“steering”* the low E-field region by rotating the coil around patient’s head, there is no other control on the shape and extend of this low field region. It is established that the SAR amplification at the tips of elongated implants depends on the coupling between the tangential component of the incident electric field and implanted wires [34, 35, 50]. As implanted DBS leads have complex trajectories consisting of segments that cannot be contained in one plane (see Figure *3*), it is important to assess the performance of any SAR-reducing strategy in models based on real patient data. This work presents the first report of such assessment, demonstrating promising results in possibility of reducing the SAR at the tips of both left and right DBS leads to levels below the SAR generated by standard body coils. It should be noted however, that the current study is limited to the assessment of heating at the tips of DBS leads in isolation, i.e., prior to their connection to the extension cables and the implanted pulse generator; thus the results presented here should not be the extended to other configurations. Further investigation is necessary to establish the efficacy of the technique in a fully implanted system.

From Table 1 it can be observed that for patients 5, 7 and 8 the optimum coil rotation angle that minimized the SAR at the tips of right and left leads was the same. A closer look at the lead trajectory in these patients (Figure 1) reveals that ipsilateral and contralateral lead trajectories are routed such that their trajectories are substantially parallel in this case. We recently showed that it is possible to implement simulation-driven instructions in the implantation of extracranial portion of DBS leads during the surgery to reduce the SAR, without requiring additional equipment or adding to the surgical time [51]. Therefore, surgeons can be instructed to subcutaneously rout the leads such that their trajectories be maximally parallel. This will allow the reconfigurable coil system to perform as efficiently in the case of patients with bilateral leads as it does for unilateral implants.

## Acknowledgment

This work was supported by the NIH grants R03EB024705 and R00EB021320.

